# Test-retest reproducibility of *in vivo* magnetization transfer ratio and saturation index in mice at 9.4 Tesla

**DOI:** 10.1101/2021.12.10.472129

**Authors:** Naila Rahman, Jordan Ramnarine, Kathy Xu, Arthur Brown, Corey A. Baron

**Affiliations:** Centre for Functional and Metabolic Mapping (CFMM), Robarts Research Institute, University of Western Ontario, London, Ontario, Canada; Department of Medical Biophysics, Schulich School of Medicine and Dentistry, University of Western Ontario, London, Ontario, Canada; Translational Neuroscience Group, Robarts Research Institute, Schulich School of Medicine and Dentistry, University of Western Ontario, London, Ontario, Canada; Department of Anatomy and Cell Biology, University of Western Ontario, London, Ontario, Canada

**Keywords:** magnetization transfer ratio, magnetization transfer saturation, reproducibility, preclinical rodent imaging

## Abstract

**Background:** Magnetization transfer saturation (MTsat) imaging was developed to reduce T1 dependence and improve specificity to myelin compared to the widely used MT ratio (MTR), while maintaining a feasible scan time. Knowledge of MTsat reproducibility is necessary to apply MTsat in preclinical neuroimaging.

**Purpose:** To assess the test-retest reproducibility of MTR and MTsat in the mouse brain at 9.4 T and calculate sample sizes required to detect various effect sizes.

**Study Type:** Prospective

**Animal Model:** C57Bl/6 Mouse Model (6 females and 6 males, aged 12 – 14 weeks)

**Field Strength/Sequence:** Magnetization Transfer Imaging at 9.4 T

**Assessment:** All mice were scanned at two timepoints (5 days apart). MTR and MTsat maps were analyzed using mean region-of-interest (ROI), and whole brain voxel-wise analysis.

**Statistical Tests:** Bland-Altman plots assessed biases between test and retest measurements. Test-retest reproducibility was evaluated via between and within-subject coefficients of variation (CV). Sample sizes required were calculated (at a 95 % significance level and power of 80 %), given various minimum detectable effect sizes, using both between and within-subject approaches.

**Results:** Bland-Altman plots showed negligible biases between test and retest sessions. ROI-based and voxel-wise CVs revealed high reproducibility for both MTR (ROI: CVs < 8 %) and MTsat (ROI: CVs < 10 %). With a sample size of 6, changes on the order of 15% can be detected in MTR and MTsat, both between and within subjects, while smaller changes (6 – 8 %) require sample sizes of 10 – 15 for MTR, and 15 – 20 for MTsat.

**Data Conclusion:** MTsat exhibits comparable reproducibility to MTR, while providing sensitivity to myelin with less T1 dependence than MTR. Our findings suggest both MTR and MTsat can detect moderate changes, common in pathologies, with feasible preclinical sample sizes.

## INTRODUCTION

Magnetization transfer (MT) imaging has been used extensively to investigate changes in myelin content and integrity in brain development, injury and white matter diseases, most notably in multiple sclerosis patients.^1,2^ MT imaging applications include both conventional contrast-weighted protocols (such as magnetization transfer ratio, MTR) and quantitative MT (qMT) methods.^3^

MT is a physical process by which macromolecules and their closely associated water molecules cross-relax with protons in the free water pool.^4^ Based on this phenomenon, it is possible to quantify the protons bound to large molecules, which are not MR visible, due to their extremely short transverse relaxation time (T2).^5^ MT contrast can be generated by applying an off-resonance radiofrequency pre-pulse (MT pulse) to selectively saturate the spectrally broad macromolecular proton pool. This saturation then transfers to the free water proton pool via MT, resulting in a decrease in the observed free water signal. The magnitude of the MT effect can be characterized by the magnetization transfer ratio (MTR):

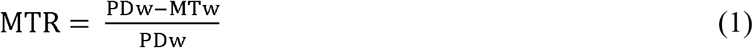

where PDw is the signal without an MT pulse applied, which is proton density weighted (PDw), and MTw is the signal with the MT pulse applied, which is MT weighted (MTw). Although MTR has been shown to correlate well with histological myelin content,^1,6^ it is also sensitive to the choice of sequence parameters, flip angle inhomogeneities, and longitudinal relaxation time (T1)^7^ T1 also correlates strongly with myelin content, but is also sensitive to axon size^8^ and iron content,^9^ mitigating the power of MTR as a measure of myelin. Quantitative MT has been used in many recent works to quantity myelin,^10–14^ as it reduces the confounding effects of scan parameters and quantifies specific tissue characteristics, such as the macromolecular pool size.^5^ However, qMT relies on complex modeling of the MR signal dependence on myelin, and requires more measurements and thus longer acquisition times compared to contrast-weighted MT protocols.^15^

Magnetization transfer saturation (MTsat) imaging was developed to improve MTR, by decoupling MTR from T1 effects, while maintaining a feasible scan time.^7^ The shorter scan time compared to qMT enables longitudinal *in vivo* imaging and allows the addition of other imaging techniques required to characterize microstructure. A scalar map of MTsat can be acquired using two reference scans of proton density and T1 weighting (PDw and T1w respectively), and one MTw scan. MTsat, being more independent of system parameters and T1 weighting, and less susceptible to inhomogeneities of the receiver coil and the transmitted RF field, provides greater specificity and contrast compared to MTR.^7,16^ MTsat shows higher white matter contrast in the brain than MTR,^7^ and has been shown to correlate more with disability metrics than MTR in patients with multiple sclerosis^17^. Hagiwara et al. reported that MTsat may be more suited to measure myelin in the white matter, compared to the ratio of T1-weighted to T2-weighted images, which has also been proposed as a measure of myelin.^18^

There is strong interest in applying MT to preclinical rodent neuroimaging studies at ultra-high field strengths, demonstrated by MTR^19–22^ and qMT studies.^13,23–25^ The feasibility of MTsat in mice at 9.4 T has been shown previously^26^ and MTsat has been explored in a feline model of demyelination at 3 T.^27^ Although most MTsat studies have been performed at 3 T, recently, Olsson et al. reported an optimized whole-brain MTsat protocol at 7 T,^28^ which highlights the increasing interest in this method. *In vivo* studies of MTsat in humans at 3 T have shown high reproducibility.^29,30^ However, to our knowledge, there are no test-retest reproducibility studies of MTsat or applications of MTsat to animal models of disease/injury at ultra-high field strength. Moreover, there are no studies comparing MTR and MTsat reproducibility. As MTsat provides a time-efficient alternative to fully quantitative techniques but with increased specificity and contrast compared to MTR, investigation of MTsat in preclinical rodent imaging will likely be of interest to other research groups. The aim of this work was to assess test-retest reproducibility of *in vivo* MTR and MTsat in mice at 9.4 Tesla and provide estimates of required sample sizes, which is essential in planning preclinical neuroimaging studies involving models of disease/injury.

## METHODS

### Subjects

All animal procedures were approved by the University of Western Ontario Animal Use Subcommittee and were consistent with guidelines established by the Canadian Council on Animal Care. Twelve adult C57Bl/6 mice (six males and six females) were scanned twice 5 days apart. The sample size was chosen to reflect similar sample sizes used in other pre-clinical imaging studies.^31–34^ Before scanning, anesthesia was induced by placing the animals in an induction chamber with 4 % isoflurane and an oxygen flow rate of 1.5 L/min. Following induction, isoflurane was maintained during the imaging session at 1.8 % with an oxygen flow rate of 1.5 L/min through a custom-built nose cone. The mouse head was fixed in place using ear bars and a bite bar to prevent head motion. These mice were part of a longitudinal study, at the end of which they were euthanized for histology. The mice were anesthetized with ketamine/xylazine (2:1) and then underwent trans-cardiac perfusion with ice-cold saline, followed by 4% paraformaldehyde in phosphate-buffer saline (PBS).

### *In vivo* MRI

*In vivo* magnetic resonance imaging (MRI) experiments were performed on a 9.4 Tesla (T) Bruker small animal scanner equipped with a gradient coil set of 1 T/m strength (slew rate = 4100 T/m/s). A single channel transceive surface coil (20 mm × 25 mm), built in-house, was fixed in place directly above the mouse head to maximize signal-to-noise ratio (SNR). A boost in SNR in the cortex when using this surface coil, compared to a commercially available 40-mm millipede (MP40) volume coil (Agilent, Palo Alto, CA, USA), has been reported previously.^35^

The MT protocol required 50 minutes total scan time and comprised three FLASH-3D (fast low angle shot) scans, and one RF transmit field (B1) map scan to correct for local variations in flip angle. An MT-weighted scan, and reference T1-weighted and PD-weighted scans (MTw, T1w, and PDw respectively) were acquired by appropriate choice of the repetition time (TR) and the flip angle (α): TR/α = 8.5 ms/20° for the T1w scan and 25 ms/9° for the PDw and the MTw scans. MT-weighting was achieved by applying an off-resonance Gaussian-shaped RF pulse (12 ms duration, 385° nominal flip angle, 3.5 kHz frequency offset from water resonance, 5 μT RF peak amplitude) prior to the excitation. Other acquisition parameters were: TE = 2.75 ms; resolution = 150×150×400 μm^3^; field of view (FOV) = 19.2 × 14.4 × 12 mm^3^; read-out bandwidth = 125 kHz; 12 averages. The B1 map was acquired acquired based on the actual flip angle imaging (AFI) method^36^ at a lower resolution of 600×600×400 um^3^ and the following scan parameters: TE = 4 ms; α = 60°; short TR = 20 ms; long TR = 100 ms; 2 averages. Anatomical images were also acquired for each subject within each session using a 2D T2-weighted TurboRARE pulse sequence (150 μm in-plane resolution; 500 μm slice thickness; TE/TR = 40/5000 ms; 16 averages; total acquisition time = 22 min).

### Image Processing

MTR and MTsat maps were generated using in-house MATLAB code. Gaussian filtering (full-width-half-maximum = 3 voxels) was first applied to the original images (MTw, PDw, and T1w images, and B1 maps) to reduce noise, while retaining image contrast. The standard MTR maps were calculated using Equation (1). MTw, PDw, and T1w images were used to calculate MTsat maps, following the original method proposed by Helms et al.,^7^ and outlined by Hagiwara et al. MTsat is inherently robust against differences in relaxation rates and inhomogeneities of RF transmit and receive field compared with conventional MTR imaging.^7,16^ Furthermore, B1 maps were used to correct for small residual higher-order dependencies of the MT saturation on the local RF transmit field to further improve spatial uniformity, as suggested by Weiskopf et al.^29^:

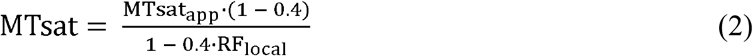

where RF_local_ is the relative flip angle α compared to the nominal flip angle, and MTsat_app_ is the uncorrected apparent MTsat. RF_local_ was calculated based on the AFI method.^36^

Brain masks were produced using the skull stripping tool from BrainSuite (v. 19b).^37^ Image registration was performed using affine and symmetric diffeomorphic transforms with ANTs software (https://github.com/ANTsX/ANTs).^38^ Region-of-interest (ROI) masks were acquired from the labeled Allen Mouse Brain Atlas.^39^ Since registration to an atlas is time-consuming, only one anatomical T2-weighted scan was chosen (the “chosen T2”) to be registered to the atlas. All other anatomical T2-weighted images were registered to the chosen T2. MTR parameter maps were registered to the corresponding anatomical images (from the same subject at the same timepoint). For ROI-based analysis, the inverse transforms resulting from these three registration steps (MTR ➔ corresponding T2 ➔ chosen T2 ➔ atlas) were then used to bring the labeled atlas to the corresponding MT space for each subject at each timepoint. Binary masks for each ROI were generated by thresholding the labeled atlas. Each mask was eroded by one voxel, except for the corpus callosum masks, to minimize partial volume errors within a given ROI. The binary masks were visually inspected to ensure good registration quality.

Furthermore, to perform whole brain voxel-wise analysis of all subjects across both timepoints, the data was registered to a common template. MTR maps were first registered to one MTR map (the “chosen MTR”). All MTsat maps were then registered to the chosen MT space using a single transform: MTR ➔ chosen MTR. For voxel-wise analysis targeted to specific ROIs, the labeled atlas was registered to the chosen MT space.

### Data Availability

The test-retest dataset and in-house code to compute MTR and MTsat is available online.^40^

### Data Analysis

#### ROI-based and Voxel-wise Analysis

ROI analysis was performed using two approaches: (1) analysis of unregistered data and (2) analysis of data registered to a common template. For the second approach, all MTR and MTsat maps were registered to a “chosen” MTR space, as described above.

The ROI analysis focused on five different tissue regions: corpus callosum (CC), internal capsule (IC), hippocampus (HC), cortex (CX), and thalamus (TH). For both ROI-based approaches, Bland-Altman and CV analyses were performed using the mean MTR and MTsat values from each ROI. Voxel-wise CV analysis was also performed with the registered data.

#### Statistical Analysis

Measurement reproducibility was explored for both ROI-based analysis and whole brain voxel-wise analysis, since both are common analysis techniques in neuroimaging. To mitigate partial volume errors from cerebrospinal fluid (CSF) in ROI-based analysis, voxels with MTR < 0.1 were omitted in both test and retest images. In voxel-wise analysis, voxels with MTR < 0.1, as measured on the test images, were omitted. Bland-Altman analysis was performed for the ROI-based analyses to identify any biases between test and retest measurements. For both analysis techniques, the scan-rescan reproducibility was characterized using the coefficient of variation (CV). The CV reflects both the reproducibility and variability of these metrics, as well as provides insight into necessary sample sizes and minimum detectable effect size. CVs were calculated between subjects and within subjects to quantify the between subject and within subject reproducibility, respectively. The between subject CV was calculated separately for the test and retest timepoints as the standard deviation divided by the mean value across subjects 1– 12. These two CV values were then averaged for the mean between subject CV. The within subject CV was calculated separately for each subject as the standard deviation divided by the mean of the test and retest scans. The 12 within subject CVs were then averaged to determine the mean within subject CV.

Sample size calculations were performed based on CVs from the ROI analysis of registered data. Minimum sample sizes required to detect defined biological effects, using both between and within subject approaches, were determined at a 95 % significance level (α = 0.05) and power of 80 % (1-β = 0.80). Following the procedure presented in van Belle,^41^ the between subject CVs were used to determine the sample size required per group to detect a defined biological effect between subjects in each ROI. Assuming paired t-tests, the standard deviations of the differences between test-retest mean values across subjects, were used to determine the sample size required to detect a defined biological effect within subjects in each ROI.^42^

## RESULTS

### Parameter Maps

Representative parameter maps are shown in Figure 1. MTsat is comparable to MTsat maps acquired by Boretius et al. in the mouse brain at 9.4 T^26^ and reveals slightly greater contrast than MTR between gray matter and white matter, which is noticeable when comparing the corpus callosum and internal capsule (white matter regions) to the surrounding gray matter.

**Figure 1.**
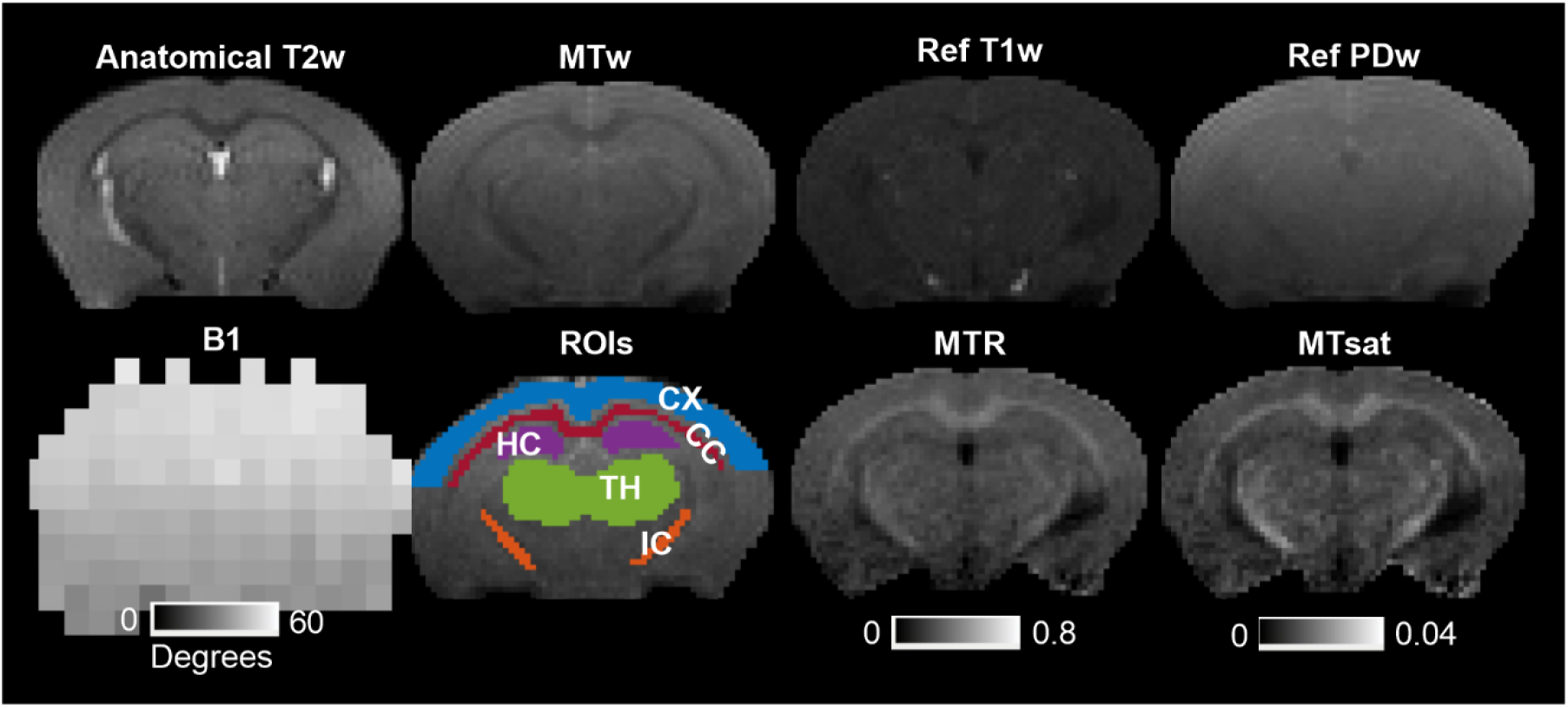
Example axial cross sections from a single subject. An anatomical T2-weighted image, an MT weighted (MTw) image, reference T1 weighted (T1w) and proton density weighted (PDw) images, a B1 map, and corresponding MTR and MTsat maps are shown. ROIs analyzed are overlaid on an MTw image and abbreviated as follows: CC – corpus callosum; IC – internal capsule; HC – hippocampus; CX – cortex; TH – thalamus.

### ROI-based Analysis

Violin plots depict the distribution of the mean values for each metric within each ROI for the 12 subjects for both registered and unregistered datasets (Figure 2). Across all metrics, the median and interquartile range are similar for test and retest conditions and results are comparable for registered and unregistered datasets. In general, the smaller ROIs (i.e., the internal capsule) showed greater distributions, while the larger ROIs (i.e., the cortex) showed much tighter distributions.

**Figure 2.**
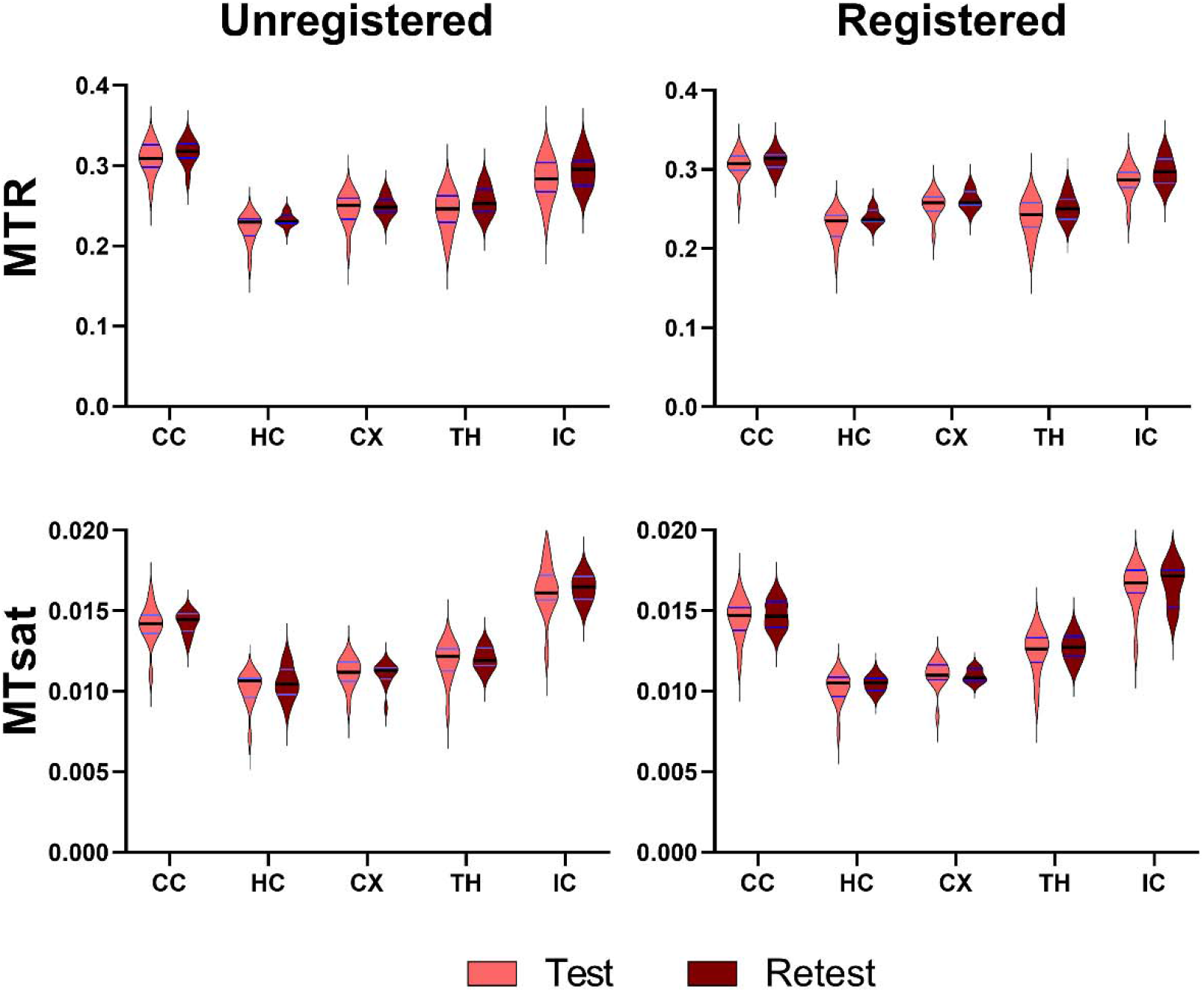
Violin plots showing the distribution of MTR and MTsat at the test and retest timepoints (five days apart) for 12 subjects in several brain regions. Unregistered data (left column) and data registered to a common template (right column) are shown. The dark black line represents the median and the red lines depict the interquartile range (25^th^ to 75^th^ percentile). The violin plots extend to the minimum and maximum values of each metric. ROIs are abbreviated as follows: CC – corpus callosum; IC – internal capsule; HC – hippocampus; CX – cortex; TH – thalamus.

Bland-Altman (BA) plots revealed negligible biases for MTR and MTsat (Figure 3). MTR exhibited lower between and within subject CVs (4.5 – 8 %) compared to MTsat (6 – 10 %), as shown in Figure 4. In general, CVs are comparable across all ROIs. These trends are comparable across both registered and unregistered data.

**Figure 3.**
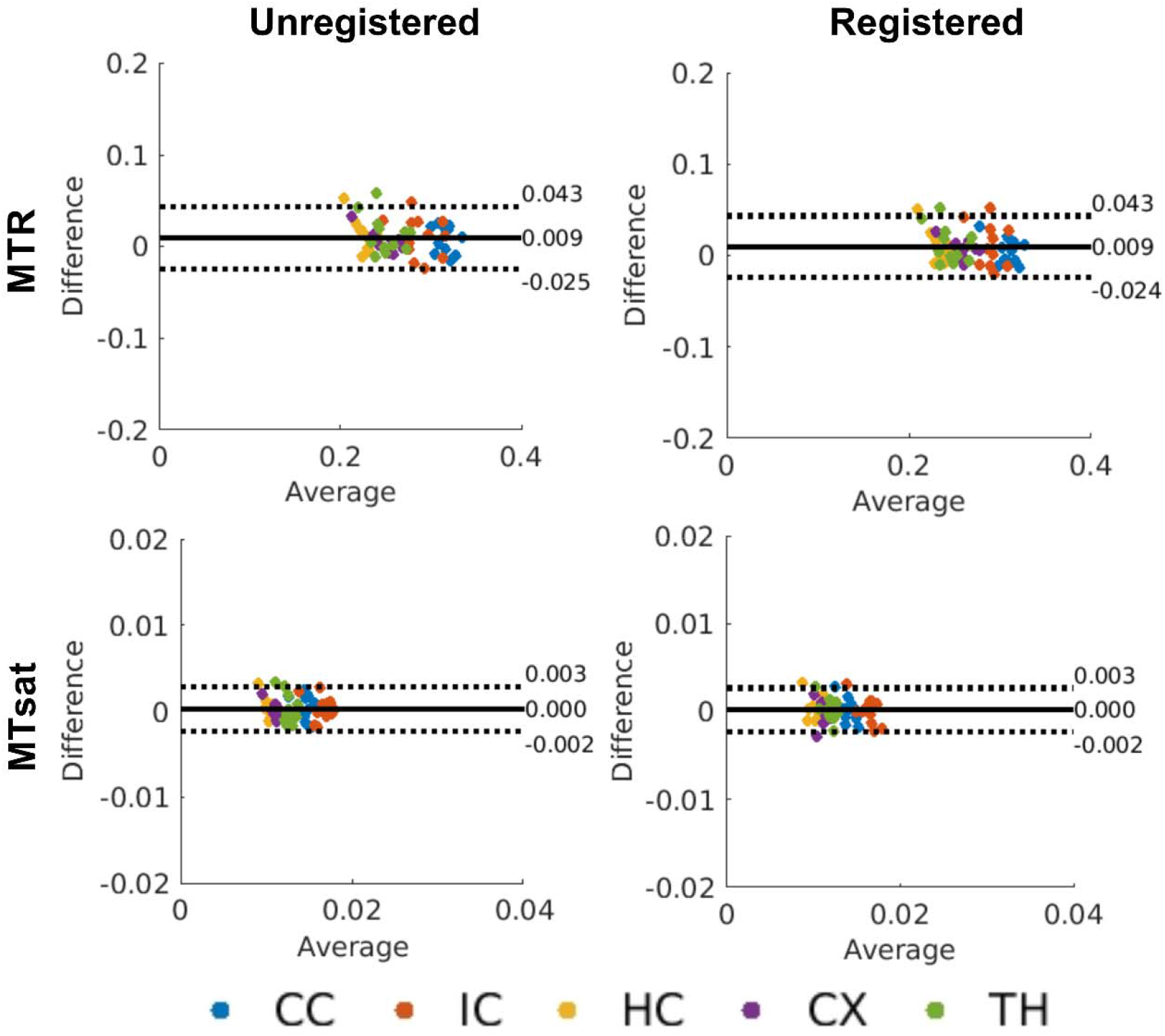
Bland-Altman plots depicting biases between test and retest scans for mean MTR and MTsat values (from the ROI-based analysis). Unregistered data (left column) and data registered to a common template (right column) are shown. The solid black lines represent the mean bias, and the dotted black lines represent the ±1.96 standard deviation lines. The average of the test and retest mean values is plotted along the x-axis and the difference between the test and retest mean values is plotted along the y-axis. ROIs in the legend are abbreviated as follows: CC – corpus callosum; IC – internal capsule; HC – hippocampus; CX – cortex; TH – thalamus.

**Figure 4.**
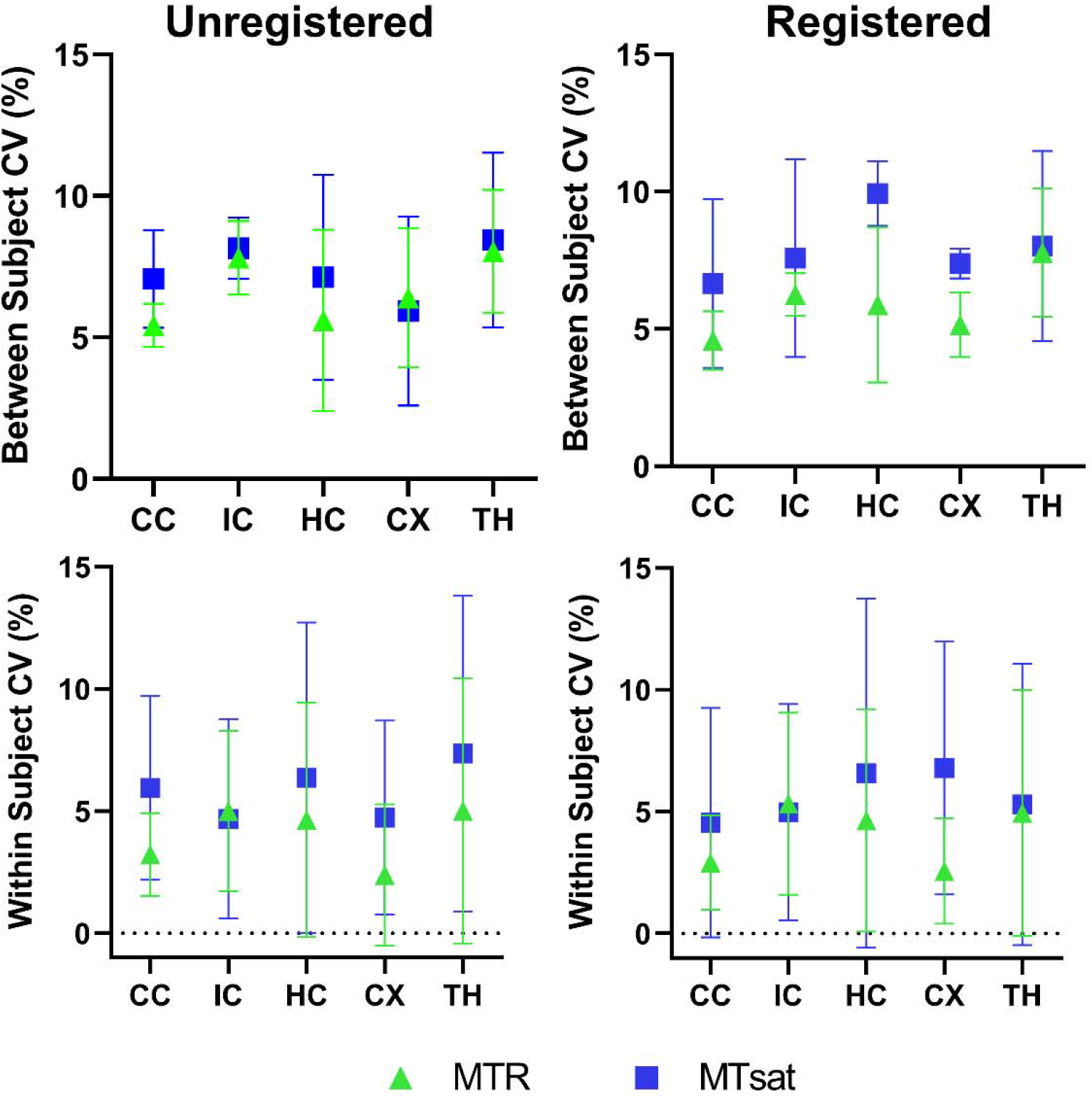
Mean between subject and within subject coefficients of variation (CV) for MTR and MTsat in each ROI. Reproducibility metrics for unregistered data (left column) and data registered to a common template (right column) are shown. Values for the between subject CV condition represent the mean ± standard deviation over subjects (averaged over the test and retest timepoints). Values for the within subject CV condition represent the mean ± standard deviation between test and retest (averaged over the 12 subjects). ROIs are abbreviated as follows: CC – corpus callosum; IC – internal capsule; HC – hippocampus; CX – cortex; TH – thalamus.

### Voxel-wise Analysis

The voxel-wise CV maps show very high CVs in the cerebrospinal fluid (CSF), due to the low values of MTR and MTsat in the CSF (Figure 5). Between and within subject CVs are stable throughout the whole brain, with the within subject CVs being slightly lower than the between subject CVs. As observed in the ROI-based CVs, MTR exhibited lower between and within subject CVs (with peaks at 7 % and 6 %, respectively) compared to MTsat (with peaks at 15 % and 12 %, respectively), as shown in whole brain histograms (Figure 6). The MTsat histograms also revealed a wider distribution compared to the MTR histograms.

**Figure 5.**
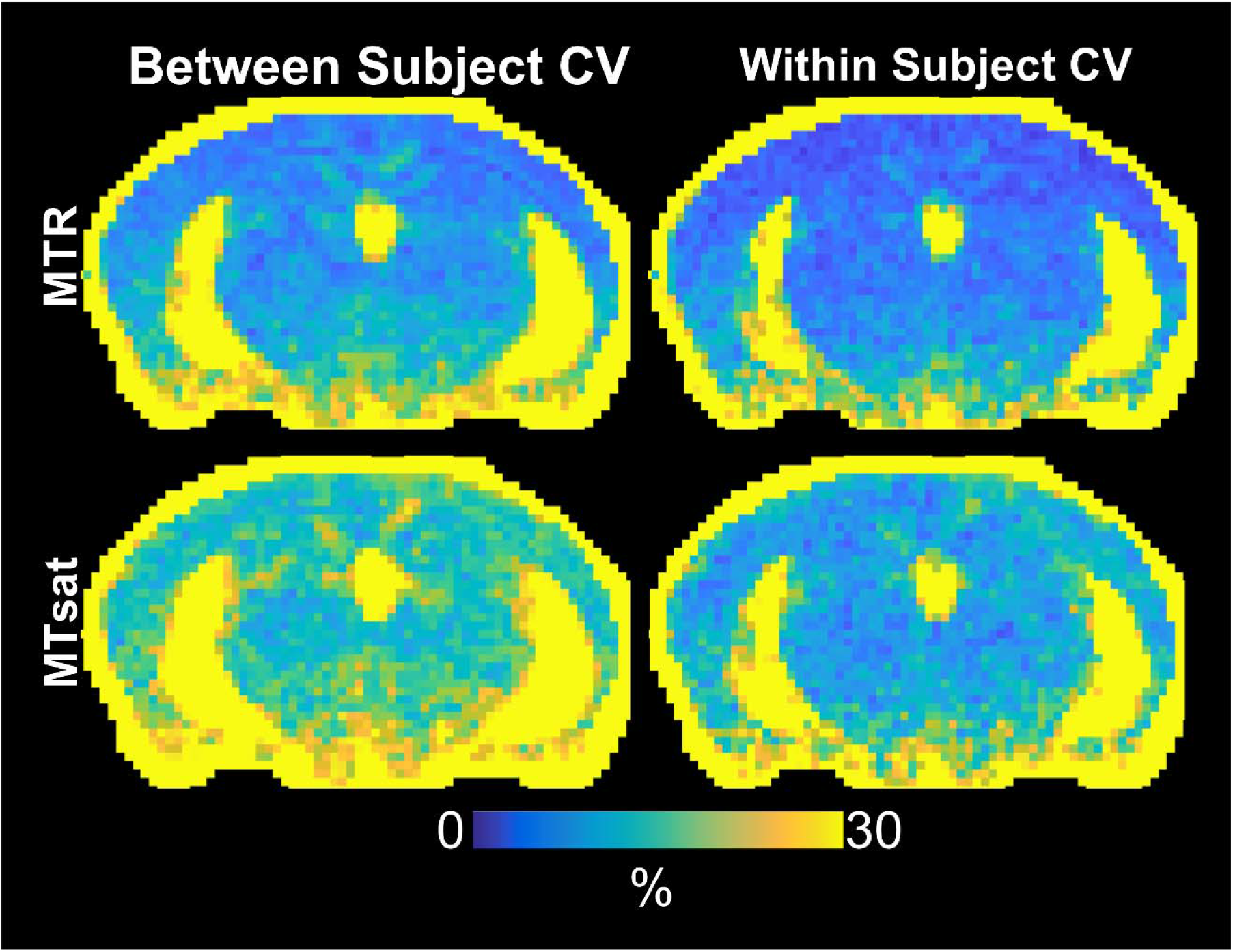
Voxelwise average between subject and within subject CV maps for MTR (top row) and MTsat (bottom row). Values for the between subject condition represent the mean CV within each voxel averaged over the test and retest timepoints. Values for the within subject condition represent the mean CV within each voxel averaged over all 12 subjects. ROIs are abbreviated as follows: CC – corpus callosum; IC – internal capsule; HC – hippocampus; CX – cortex; TH – thalamus.

**Figure 6.**
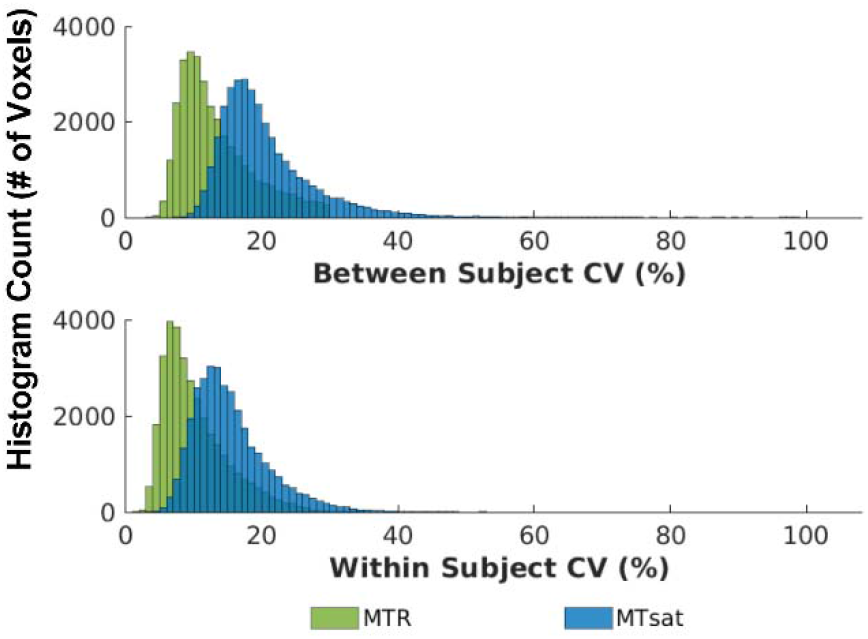
Distribution of whole brain voxel-wise between and within subject CVs for MTR and MTsat.

### Sample sizes and minimum detectable effect

#### Between subjects

The between subject CVs, from the ROI analysis, were used to determine the minimum sample sizes required to detect statistically significant changes of 6, 8, 10, 12 and 14 % between subjects in each metric within each ROI. To detect a minimum change of 8 % in all ROIs, MTR required a sample size of 15 (Figure 7). In comparison, MTsat required a sample size of 25 to detect an 8 % change in all ROIs. The CC and CX required smaller sample sizes, with MTR requiring 12 subjects to detect a 6 % change, and MTsat requiring 15 subjects to detect an 8 % change.

**Figure 7.**
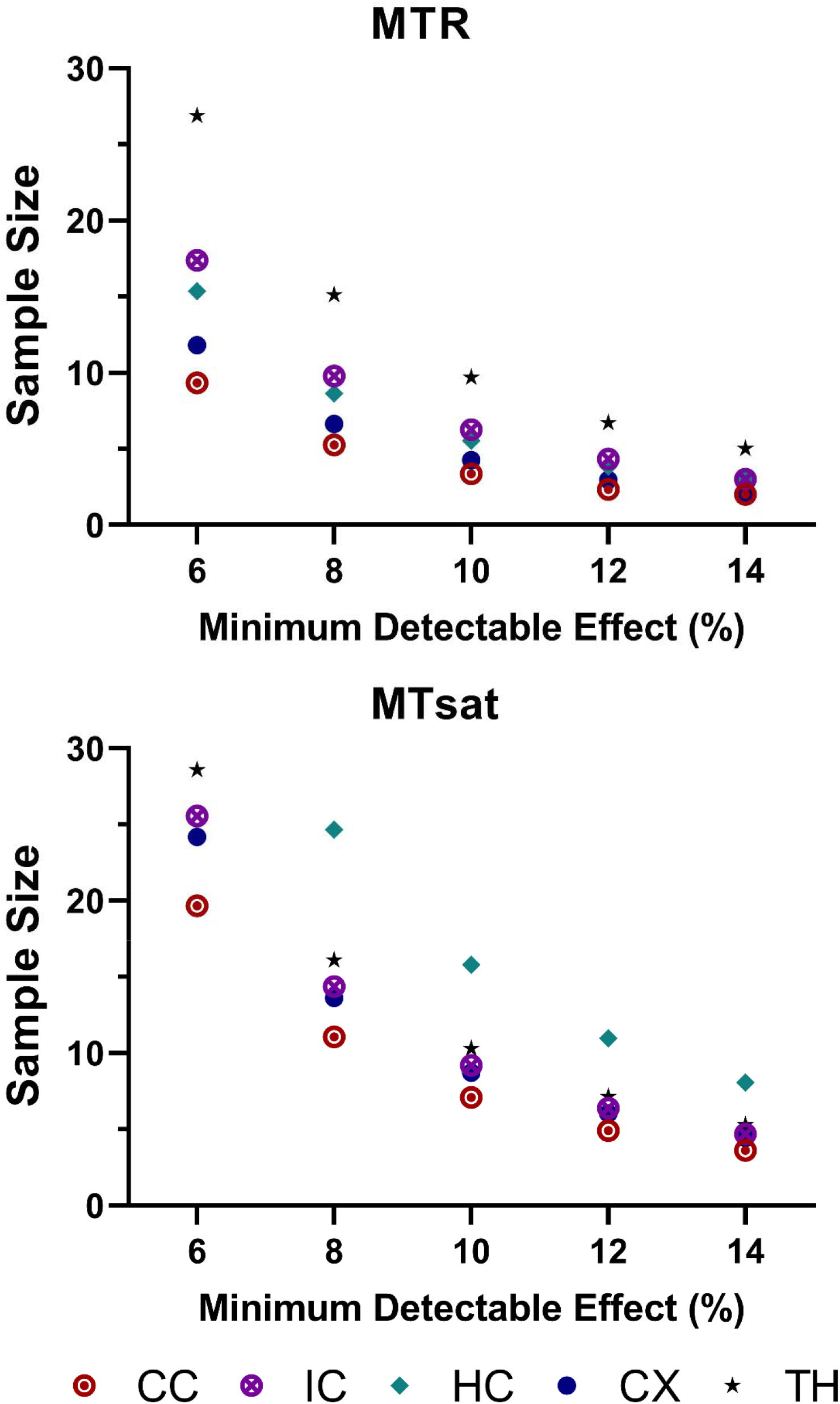
Sample size estimation using a between-subjects approach on data registered to a common template. Sample sizes required, calculated from ROI-based between-subject CVs, to detect a statistically significant effect within each ROI with a change in each metric of 6, 8, 10, 12, and 14 %. Note that the sample size range varies between plots and sample sizes exceeding the range are not shown. ROIs are abbreviated as follows: CC – corpus callosum; IC – internal capsule; HC – hippocampus; CX – cortex; TH – thalamus.

**Figure 8.**
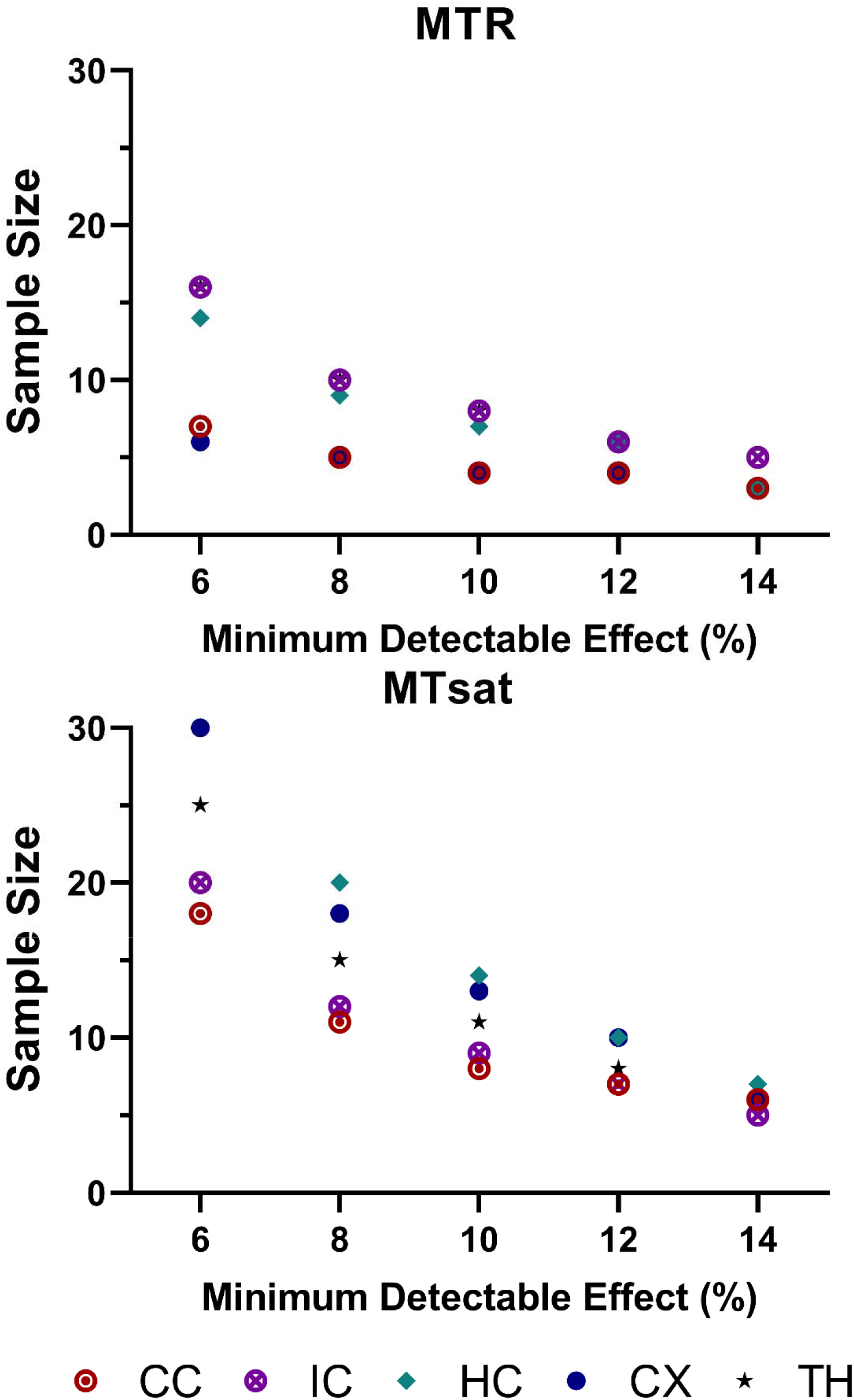
Sample size estimation using a within-subjects approach on data registered to a common template. Sample sizes required, calculated from the standard deviation of differences between test-retest mean values across subjects (assuming paired t-tests), to detect a statistically significant effect within each ROI with a change in each metric of 6, 8, 10, 12, and 14 %. Note that the sample size range varies between plots and sample sizes exceeding the range are not shown. ROIs are abbreviated as follows: CC – corpus callosum; IC – internal capsule; HC – hippocampus; CX – cortex; TH – thalamus.

#### Within subjects

The standard deviations of the differences between test-retest mean values across subjects (assuming paired t-tests) were used to determine the minimum sample sizes required to detect statistically significant changes of 6, 8, 10, 12, and 14 % within subjects in each metric within each ROI. In the CC and CX, minor changes (6 %) can be detected in MTR with 6 subjects per group, while MTsat could detect larger changes (8 – 12 %) with 12 subjects per group. For MTR, small changes (6 %) could be detected in the other ROIs (IC, HC, TH) with a feasible sample size of 15. MTsat can detect larger changes (8 % and greater) in all ROIs with 20 subjects per group.

## DISCUSSION

This study explores the reproducibility of MTR and MTsat at 9.4 T. No biases were found between repeat measurements with ROI-based analysis. MTR and MTsat were shown to be reproducible in both the mean ROI analysis and the whole brain voxel-wise analysis, with MTsat CVs being slightly higher than MTR CVs. Overall, within subject CVs were lower than between subject CVs for both ROI-based and voxel-wise analysis, indicating less variability within subjects on a test-retest basis.

### ROI-based Reproducibility

ROI-based reproducibility was investigated using an unregistered dataset and a dataset registered to a common template, as both unregistered and registered analysis techniques have been used in neuroimaging studies, and the difference between using either analysis technique remains sparsely explored. Recently, Klingenberg et al. reported that registration significantly increased the accuracy of a convolutional neural network (CNN) to detect Alzheimer’s disease, compared to no registration.^43^ Violin plots, BA plots, and ROI-based CV analysis reveal the same trends for both registered and unregistered ROI-based analysis approaches, which indicates that either method can be used for MT analysis. However, we recommend using the registered analysis approach, as there is only one set of ROI masks to edit, making the analysis process more time efficient. The unregistered analysis approach will also introduce inter- and intra-rater variability, due to the large number of ROI masks being edited.

The MTR ROI CVs observed in this work are consistent with MTR CVs in human studies done by Welsch et al.^44^ and Hannila et al.^45^ MTsat CVs reported here are comparable to MTsat CVs in human studies at 3 T.^29,30^ Overall, MTsat exhibits slightly higher CVs than MTR, which may arise from noise propagation through the equations used to calculate MTsat, as described by Olsson et al.^28^ A noticeable increase in MTsat CVs compared to MTR CVs, in the HC and CX, may be due to low MTsat values in these regions.

### Voxel-wise Reproducibility

Voxel-wise CV trends were comparable to ROI-based CV trends. Voxel-wise CV maps revealed a more noticeable increase in CVs in the superior-inferior direction of the brain in MTR, compared to MTsat. This can be related to the inherent compensation of flip angle inhomogeneities in MTsat.

### Sample Size and Minimum Detectable Effect

Given the current test-retest study design, small changes (6 - 8 %) can be detected in MTR and MTsat, both between and within subjects, with feasible sample sizes of 10 – 15 for MTR, and 15 – 20 for MTsat. With a sample size of 6, moderate changes (~15 %) can be detected in MTR and MTsat, both between and within subjects, in all ROIs. The CC consistently exhibited the smallest required sample sizes, which can be related to the lower variability of myelin content in the CC, compared to the gray matter ROIs.^46^ Interestingly, the CC and IC (the white matter regions) require similar sample sizes to detect the same changes in MTsat (using both between and within subject approaches), but not in MTR, which requires larger sample sizes to detect changes in IC. This may stem from the better contrast seen between the IC and gray matter in MTsat, compared to MTR, which arises from MTsat being less susceptible to inhomogeneities of the transmitted field and more independent of T1-weighting.^7,26^ Most MT studies report changes in MTR between 15 – 30 %, with some studies reporting more subtle changes between 5 – 10 %. In a cuprizone demyelination model in mice, MTR decreased by 15 % and 30 % at 4 weeks and 6 weeks of cuprizone administration, respectively.^47^ In an ischemic injury model in mice, MTR decreased by 30 % in the corpus callosum of injured mice compared to controls.^21^ In a closed head traumatic brain injury model in mice, MTR in the corpus callosum decreased by 10 % from baseline at 1-day post-injury.^20^ A post-mortem study revealed a 10 % decrease in MTR between normal-appearing white matter and multiple sclerosis lesions.^1^ In a recent multiple sclerosis study, MTR was able to differentiate between patients with and without cognitive impairment, showing a 7 % decrease in patients with cognitive impairment.^48^

MTR can detect changes on the order of 15 - 30 % (such as the changes found in the cuprizone demyelination model) with small sample sizes (n = 6). With disease and injury models resulting in less drastic changes to myelin content, our findings suggest that MTR and MTsat can detect smaller changes with feasible preclinical sample sizes. Thiessen et al.^47^ showed that when there’s an 80% reduction in myelinated axon density, MTR only decreases by ~ 30 % (because it’s thought that inflammation has a competing effect on MTR). So, a two-fold difference in myelination will result in at least a 15 % change in MTR. However, as MTsat provides greater specificity to myelin, a two-fold difference in myelination should translate to a larger percent change in MTsat.

### Limitations

Although a volume coil is more appropriate for structural imaging as it provides stable signal-to-noise ratio throughout the brain, this study used a transceive surface coil. The voxelwise CV maps show that between-subject and within-subject CVs are slightly higher towards the inferior region of the brain. However, the increase in CV is subtle and as shown in ROI-based analysis, the CVs of ROIs located in inferior regions of the brain (such as the IC) are comparable to the ROIs closer to the surface coil. This shows the feasibility of acquiring MTR and MTsat data using a surface coil, which may be useful in studies in which MT imaging is combined with other methods that require a surface coil or inherently low SNR methods that would benefit from a surface coil, such as diffusion MRI. Recent preclinical investigations have included a combination of MT imaging and diffusion MRI.^20,25,49^ Moreover, the findings in this study will complement a recent test-retest reproducibility study in advanced diffusion MRI techniques in mice at 9.4 T.^35^

In the statistical analyses, it should be noted that for the within-subject calculation of CV, the standard deviation was determined from only two data points (the test and retest conditions). As a result, the standard deviation may not accurately represent the spread of data within the population, leading to an unknown bias in the resulting within-subject CV.

## CONCLUSION

In conclusion, we have investigated the reproducibility of MTR and MTsat in a rodent model at an ultra-high field strength. We have shown that MTR and MTsat are reproducible in both ROI-based analysis, which includes both registered and unregistered analysis techniques, and voxel-wise analysis. Importantly, MTsat exhibits comparable reproducibility to MTR, while providing better contrast. With a sample size of 6, changes on the order of 15% can be detected in MTR and MTsat, both between and within subjects, while smaller changes (6 – 8 %) require feasible sample sizes of 10 – 15 for MTR, and 15 – 20 for MTsat. This work will provide insight into experiment design and sample size estimation for future longitudinal *in vivo* MTsat imaging studies at 9.4 T.

## Notes

<u>Grant Support:</u> The authors would like to acknowledge the Canada First Research Excellence Fund (BrainsCAN - https://brainscan.uwo.ca/); the New Frontiers in Research Fund (NFRFE-2018-01290 - https://www.sshrc-crsh.gc.ca/funding-financement/nfrf-fnfr/index-eng.aspx), awarded to CB; the Natural Sciences and Engineering Research Council of Canada: Canada Graduate Scholarships - Master’s Program (NSERC-CGS M), awarded to NR; and the Ontario Graduate Scholarship (OGS), awarded to NR, for their funding contributions.

https://osf.io/5nwae/

### Competing Interest Statement

The authors have declared no competing interest.

